# An experimental test of the mutation-selection balance model for the maintenance of genetic variance in fitness components

**DOI:** 10.1101/193425

**Authors:** Nathaniel P. Sharp, Aneil F. Agrawal

**Author notes:** Corresponding author: Nathaniel Sharp, Department of Zoology, University of British Columbia, #4200-6270 University Blvd., Vancouver, BC Canada V6T 1Z4.Ph. Author contributions: NS and AA designed experiments; NS performed experiments; NS and AA analyzed the data and wrote the manuscript.

## Abstract

Despite decades of research, the factors that maintain genetic variation for fitness are poorly understood. Mutation selection balance will always contribute to standing variance, but it is unclear what fraction of the variance in a typical fitness component can be explained by mutation-selection balance and whether fitness components differ in this respect. In theory, the level of standing variance in fitness due to mutation-selection balance can be predicted using the rate of fitness decline under mutation accumulation, and this prediction can be directly compared to the actual standing variance observed. This approach allows for controlled statistical tests of the sufficiency of the mutation-selection balance model, and could be used to identify traits or populations where genetic variance is maintained by factors beyond mutation-selection balance. For example, some traits may be influenced by sexually antagonistic balancing selection, resulting in an excess of standing variance beyond that generated by deleterious mutations. To encourage the application of this approach, we describe the underlying theory and use it to test the mutation selection balance model for three traits in *Drosophila melanogaster*. We find some evidence for non-mutational variance in male mating success and female fecundity relative to larval viability, which is consistent with balancing selection on sexual fitness components. Finally, we discuss the theoretical and practical limitations to this approach, and discuss how to apply it successfully.

## INTRODUCTION

The maintenance of genetic variation for fitness has been called one of the most important unresolved issues in evolutionary biology (Lynch and Walsh 1998; Barton and Keightley 2002). Despite the action of natural selection, populations are found to harbour significant genetic variation for fitness-related traits (Houle 1992). A number of possible sources of variation exist, including deleterious mutations, beneficial alleles on their way to fixation, and balancing selection, including environmental heterogeneity in selection. The relative importance of these factors is unknown, and will determine how genomes and populations evolve (Mitchell Olds *et al.* 2007).

Deleterious mutation-selection balance is arguably the most general explanation for genetic variance in fitness because all populations experience mutation. The question is not whether mutation-selection balance contributes to standing variation but rather whether it alone accounts for most or all of the variation. Answering this question requires knowing how much variation would be expected from mutation selection balance alone. In a population at equilibrium, the rate at which mutation reduces mean fitness will equal the rate at which selection increases mean fitness, which in turn is equal to the genetic variance in fitness (Fisher 1930). Thus, under mutation-selection balance, the standing genetic variance in fitness should be equal to the rate of fitness decline per generation in a mutation accumulation experiment (“mutational decline”). This idea is described more formally below, and can be found in various incarnations in the literature (reviewed in Barton 1990; Houle *et al.* 1996; Charlesworth and Charlesworth 2012; Charlesworth 2015; Charlesworth and Hughes 2000). If there is any other source of genetic variation apart from deleterious mutations, then the observed level of standing variation will exceed the value expected from deleterious mutations. Mutation-selection balance is therefore a relevant and testable hypothesis for the maintenance of variation.

Although estimates of mutational decline and standing variance exist for a number of traits (particularly in *D. melanogaster*), they are generally estimated in different experimental populations with different genetic backgrounds using different methodologies, which is far from ideal, especially given the considerable variation in some quantitative genetic parameters among populations (Houle 1998; Charlesworth 2015). In addition, the effects of new mutations are often assessed in the homozygous state; because new mutations will rarely be found in the homozygous state in a randomly-mating population, it is their heterozygous effects that are most relevant to the expected level of standing variation. Homozygous mutational decline can only be used to infer the heterozygous equivalent after making tenuous assumptions about the average dominance of mutations.

Despite these challenges, several attempts have been made to qualitatively test alternative models for the maintenance of genetic variation using data from numerous sources (Houle *et al.* 1996; Houle 1998; Charlesworth 2015; Charlesworth and Hughes 2000). These investigations point to an important role of deleterious mutation in maintaining variation, but they cannot statistically evaluate the adequacy of the mutation-selection balance model. While these studies have provided valuable insights, their conclusions are often limited by the quality and quantity of available data, leading these authors and others to call for experiments that directly address the mutation-selection balance hypothesis (Keightley 1994; Lynch *et al.* 1999; Charlesworth *et al.* 2007). Here we outline the theoretical and empirical issues relevant to such experiments, and use this approach to study the sources of standing genetic variation in *Drosophila melanogaster*.

Populations in the laboratory, which are generally maintained in constant conditions for many generations, may be unlikely to harbour certain kinds of genetic variation because spatial and temporal variation in selection will be minimal, and beneficial mutations on their way to fixation will be rare if the population is well adapted. (Moreover, there is reason to believe that beneficial alleles that eventually become fixed may make a relatively small contribution to standing variance (Charlesworth 2015).) However, although the external environment may be relatively constant, alleles in sexual populations will be expressed in both males and females, and may be subject to intra-locus sexually antagonistic (SA) selection. This type of balancing selection may be a particularly important source of variation in well-adapted populations (Long *et al.* 2012).

Although the existence of SA alleles has been established, it remains an empirical challenge to quantify their contribution to the standing genetic variance in fitness, relative to other sources of variation. If alleles under SA balancing selection are common, deleterious mutation should account for a smaller fraction of the genetic variance in sex-specific fitness components than in non-sex-specific fitness components. In other words, SA is expected to generate “excess” genetic variance beyond that attributable to deleterious mutations. To test this idea we collected data on standing variance and the rate of change due to mutation accumulation (MA) for larval viability, male mating success, and female fecundity. We studied heterozygous second chromosomes (∼37% of the genome) on a common isogenic background, and measured mutational decline and standing variance in the same way within each fitness component. We find evidence for excess variation in the sexual (adult) fitness components, which is consistent with some form of balancing selection involving these fitness components, the most obvious of which is sexual antagonism. However, the standing genetic and mutational correlations between fitness components do not present a strong signature of sexual antagonism, suggesting that other sources of non mutational variation may be present.

## THEORETICAL BACKGROUND

The theoretical background motivating this experiment has been described by others (Barton 1990; Houle *et al.* 1996; Charlesworth and Charlesworth 2012; Charlesworth 2015; Charlesworth and Hughes 2000), and we present a summary of the major points here. What follows is one of several possible approaches to the theory surrounding mutation-selection balance, and the terms used generally follow those of Charlesworth and Charlesworth (2012, pp. 183-6). We present an additive model formulation below; an analogous multiplicative formulation where traits are measured on a log scale is given in the Supporting Information.

In a large population with random mating, a deleterious mutation at a given locus will be present at a frequency of *q*\* ≈ *μ*/(*hs*), where *μ* is the mutation rate and *hs* is the coefficient of selection against heterozygotes (Haldane 1937). This classic result assumes that mutation is weak relative to selection, and that *q*\* will be very small as a result, such that the deleterious allele will almost never occur in the homozygous state, i.e., (*q\**)^2^ ≪ 1 and the homozygous mutant genotype can be ignored. Similarly, we can approximate the frequency of heterozygotes as 2*q*\*(1 – *q*\*)≈ 2*q*\*. If the non-mutant value of trait *z* is *k*, and the heterozygous trait value is *k*(1 –*a*), at equilibrium the expected trait value is 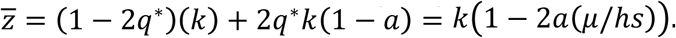 We can assume that the effect of a mutation on trait *z* is some fraction *c* of its effect on total fitness, i.e., *a* = *chs*, so that 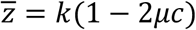. When the fraction *c* of its effect on total fitness, i.e., *a* = *chs*, so that When the trait is fitness itself, *c* = 1, and this gives the classic result that the reduction in mean fitness at mutation-selection balance relative to a mutation-free population depends only on the mutation rate (Haldane 1937).

Now consider the additive genetic variance in trait values relative to the trait mean, 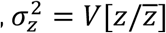 also called “evolvability” (Houle 1992). If mutation is the only source of variation, at equilibrium 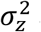 at one locus will be

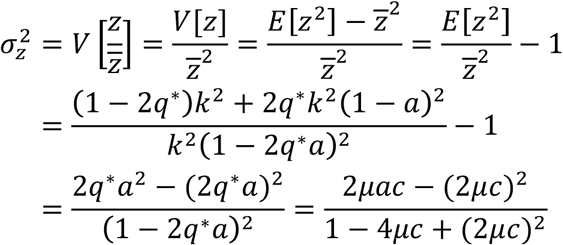

Since *μ* ≪ 1, terms in *μ*^2^ are negligible and the denominator in this relationship will be very close to 1, this leaves 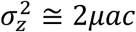. Summing over *n* loci in the genome, if the variance at each site is small and assuming there is no epistasis (Charlesworth and Hughes 2000) and no covariance between the mutation rate and the mutational effect, the total standing genetic variance in relative *z* is

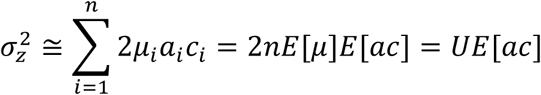

where *U* = 2*nE*[*μ*] is the mutation rate per genome. When the trait is fitness itself, *c* = 1, *a* = *hs*, and 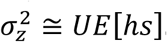. This in turn is equal to the rate of change in mean relative fitness due to one generation of mutation in the absence of selection, Δ*M*_*w*_, which can be estimated in an MA experiment (Halligan and Keightley 2009). A conceptual way to view this result is that when the effects of mutation and selection are at equilibrium the rate of decline in fitness due to mutation must equal the rate of increase in fitness due to selection, which is given by the additive genetic variance in fitness (Fisher 1930), or, for a trait, the additive genetic covariance between the trait and fitness (Price 1970). A corresponding result is that deleterious mutations contribute little in the way of dominance variance, relative to additive variance, so that most genetic variance will be additive under mutation-selection balance (Charlesworth and Charlesworth 2012, pp. 184-185). Under the null hypothesis that deleterious mutations completely account for standing variation, we expect no “excess” variance, i.e. 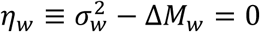. Thus *ɳ_w_*> 0 is an indication that there is more standing genetic variation than can be explained by mutation selection balance alone.

For traits that are components of fitness, standing variance will also depend on *c*, the relationship between the effect of a mutation on the trait and its effect on fitness. The standing variance in trait *z* depends on the average value of *c* across mutations, as well as the covariance between *a* and *c*, *C*[*a*, *c*]:

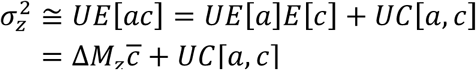

While Δ*M*_*z*_ can be estimated directly in an MA experiment, the other terms are more difficult to determine, as they depend on the pleiotropic effects of mutations on different fitness components. Observations of positive mutational correlations among fitness components (e.g., Houle *et al.* 1994; Keightley and Ohnishi 1998; Mallet *et al.* 2011; Sharp and Agrawal 2013) suggest pleiotropy is generally positive, such that the average deleterious mutation will have a smaller effect on a single fitness component than on total fitness, i.e., 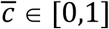, although this is not necessarily the case for all loci. Positive pleiotropy also suggests that *C*[*a*,*c*] will be positive and not too large, i.e., mutations with greater effects on a given fitness component will tend to have greater effects on total fitness. Therefore, Δ*M*_*z*_ alone would overe underestimate 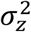 because 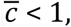 but would underestimate 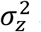 because *UC*[*a*,*c*] > 0. However, given realistic values the first bias will tend to be larger, such that Δ*M*_*z*_ will tend to overestimate 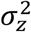, particularly when 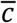 is not very close to 1, which is likely to be the case; a formal analysis of these biases is given by Charlesworth (2015). This bias means that the rate of mutational decline will tend to overestimate the expected variance in a given trait under mutation-selection balance because the frequency of mutations will be reduced through selection on additional fitness components that are not accounted for by 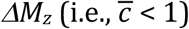. In practice, a test of the mutation selection balance hypothesis for a fitness component, mutation-selection balance hypothesis for a fitness component, 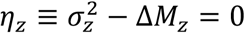 will therefore be conservative.

It may be possible to reduce this conservative bias by estimating 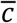. If *j* components of fitness are measured, a very simple estimate of 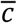 for trait *z* is the mutational decline in the trait relative to the mutational decline in all fitness components, i.e., 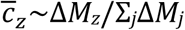 This assumes that fitness components are multiplicative but that mutational effects on individual traits are sufficiently small that the total fitness effect can be approximated via the sum across all fitness components. Even when all fitness components have been measured, this approach can over- or under-estimate 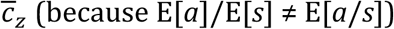. If an important fitness component has not been measured, Σ *j*Δ*M*_*j*_ < Δ*M*_*w*_, and so Δ*M*_*z*_/Σ *j*Δ*M*_*j*_ will tend to will tend to overestimate. 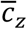. Again, this would lead to a conservative test of 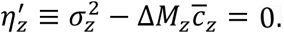 We obtained estimates of 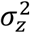 and Δ*M*_*z*_ for three fitness components (traits), as described below to test the mutation-selection balance hypothesis.

In some organisms an additional source of bias will arise if the ancestral control genome cannot be preserved, such that it is not mutation-free when measured following mutation accumulation. In other words, deleterious alleles segregating in a control population will cause mutational fitness decline, and therefore the expected variance under mutation-selection balance, to be under estimated. We address this issue further below.

## MATERIALS AND METHODS

### Overview

Our goal was to estimate the rate of mutational decline, Δ*M*_*z*_, and the standing additive genetic variance in relative trait values, 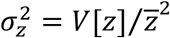, for each of three major fitness components: viability, female fecundity, and male mating success, in a simple population – fruit flies living in the lab. To do so we examined each trait in a number of mutation accumulation lines, in their corresponding controls, and in an outbred laboratory population. In each case, focal second chromosomes were tested in the heterozygous state on an isogenic background, allowing us to compare traits and sources of variance while minimizing background effects. All reported estimates refer to haploid second chromosomes. Details of line preparation and assay crosses are shown in Figures S1 and S2.

### Stocks and crosses

The outbred lab population was collected in 1970 in Dahomey (now Benin) West Africa, and maintained in the current lab for more than 3 years (>75 generations) before this experiment in a population of several thousand adults, with overlapping generations. This population and all other stocks were maintained under standard conditions (25ଌ 70% RH; 12:12 L:D). Stocks bearing visible genetic markers and balancer chromosomes, which suppress recombination on the homologous chromosome, were originally obtained from the Bloomington Drosophila Stock Center (Bloomington, IN). Except where noted, all crosses took place in 37 mL vials containing 7 mL of yeast-sugar-agar food seeded with live yeast, using virgin females where appropriate. All flies used to initiate male and female fitness assays were virgins. Experimental flies were typically 2-6 days post eclosion at the time of crossing. For both mutational decline and standing variance, all traits were measured for heterozygous focal second chromosomes situated on a common isogenic background derived from the outbred Dahomey population using standard balancer chromosome techniques. Additional stocks of this isogenic background with isogenic markers and balancer chromosomes were also created as required.

### Mutation accumulation and control lines

We measured Δ*M*_*z*_ in lines that accumulated mutations for 52 generations (2 years). The mutation accumulation (MA) procedure used to generate these lines is described elsewhere (Sharp and Agrawal 2012). Briefly, an initially-isogenic focal second chromosome marked with *bw* was maintained by crossing a single heterozygous male to outbred stock females with marker mutations for each line, thereby maximizing the amount of drift and rendering selection ineffective (Figure S1A,B). The same initial chromosome was also maintained in three separate control populations of 450 adults each. This moderate population size should limit the accumulation of new mutations. Following 52 generations of MA, crosses were performed to situate 55 MA chromosomes and 57 control chromosomes on the isogenic background described above (Figure S1C). For each of these MA and control lines, we assessed each trait in six replicates, for a total of ∼678 replicates per trait. In this MA procedure, focal chromosomes are maintained in males, and should therefore have a very low rate of recombination. However, we found evidence of recombination in four of the 55 MA lines (Sharp and Agrawal 2016). We excluded those four lines from the present study, leaving 51 MA lines. The lines used in this experiment are inferred to contain an average of 18.6 mutations each, or about 950 in total (Sharp and Agrawal 2016). In the homozygous state, these MA chromosomes cause significantly reduced viability (Sharp and Agrawal 2012) and adult reproductive fitness (Sharp and Agrawal 2013).

### Standing variance lines

We measured 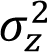 among second chromosomes derived from virgin females collected at random from the outbred Dahomey population. Crosses were performed to situate each chromosome on the isogenic background described above (Figure S2A). These crosses were designed to minimize among-line selection and to ensure that each line ultimately contained only a single focal chromosome haplotype. In addition, second chromosomes remained heterozygous throughout the experiment, which should minimize selection against genotypes bearing recessive lethal or deleterious alleles. For each of 133 such lines we assessed each trait in six replicates, for a total of ∼798 replicates per trait.

### Viability assays

Viability was estimated as the probability of survival to adulthood, allowing for larval competition with a standard genotype. In each assay replicate, two males heterozygous for the focal chromosome and an isogenic chromosome bearing the dominant phenotypic marker *L* were crossed to two females heterozygous for an isogenic standard second chromosome and an isogenic balancer chromosome, *CyO*. After four days, these adults were removed; offspring were scored up to 15 days following vial initiation. We scored the frequency of offspring bearing the focal chromosome (heterozygous with the standard isogenic chromosome) compared with the frequency of *L*/*CyO* offspring (Figure S2C). Standing variance in juvenile viability was assessed in two blocks of 67 and 66 lines respectively, and MA/control chromosomes were assessed in a separate block.

### Male mating success assays

Male mating success was measured under male-biased sex ratio conditions, in competition with males homozygous for a ubiquitously-expressed phenotypically dominant red fluorescent protein allele (*DsRed*). In each assay replicate, five virgin focal males (heterozygous for the focal chromosome), five virgin competitor males, and six virgin isogenic females were allowed to interact for 3 ± 0.25 hours (Figure S2D). This should provide sufficient time for mating to take place, while limiting the opportunity for multiple mating (Manning 1967). The males were then discarded; 1-2 days later females were placed individually into 16 mL oviposition tubes, containing ∼2 mL of food, without live yeast. After a further 8-10 days, each group of six oviposition tubes was examined to determine the number of females that produced offspring, and examined under fluorescent light to determine the number of females who produced offspring sired by *DsRed* competitor males (*DsRed* is visible in larvae, pupae, and adults). There was no evidence of multiple mating, i.e., both fluorescent and non-fluorescent offspring in a single tube, except in one case, which was removed from the dataset. Some females either died or did not produce any offspring; on average, more than five out of six females produced offspring per replicate. These assays were performed in two blocks for the MA group, with non-overlapping sets of mutant and control lines in each block (block 1: 29 control lines, 25 MA lines; block 2: 28 control lines, 26 MA lines). Similarly, two blocks were performed for the assay of standing variance in this trait, of 67 and 66 lines respectively.

### Female fecundity assays

Female fecundity was estimated as early-life egg production. Virgin focal females (heterozygous for the focal chromosome) and outbred virgin brown-eyed females (*bw*/*bw*) were held in individual vials with food but without live yeast for four days. In each assay replicate, a single focal female was then placed in a vial with a single *bw*/*bw* female and two isogenic males (Figure S2E). Each vial was supplemented with ∼10 μl of 0.02 g mL^-1^ live yeast in solution. This amount of yeast can be consumed by a single female in < 24 h (Zikovitz and Agrawal 2013), and may therefore represent a “limited” resource for females in this context, where two females are present. After 24 ± 0.5 h focal females were placed in individual oviposition vials without live yeast, containing standard media with added food coloring to facilitate egg counting. Females were allowed to oviposit for 18 ± 0.5 hours, after which they were discarded, and the number of eggs was scored. Note that the adult fitness components we measured do not involve the number of offspring produced, and are therefore independent of larval viability.

### Data analysis

We first used generalized linear mixed models fit by maximum likelihood implemented in *R* (R core Team 2014) using the *lme4* package (Bates et al. 2014) to estimate means and variance components on a log scale (see Supporting Information). The response variable for viability was the ratio of the number of offspring of the focal genotype, *n*_*focal*_, to the number of offspring of the standard genotype, *n*_*standard*_ in each replicate. When modeled using a binomial link function, the model scale logit(*n*_*focal*_/(*n*_*focal*_ + *n*_*standard*_)) is equivalent to log(*n*_*focal*_/*n*_*standard*_). The response variable for female fecundity was number of eggs in each replicate, modeled using a Poisson (log) link. The response variable for male mating success was the ratio of the number of females that mated with focal males to the number of females that mated with standard males in each replicate, modeled using a binomial link as for viability. Each trait and group of lines (MA lines, MA controls, outbred lines) was modeled separately, with a random effect of genotype (line). In addition, we included a random effect of replicate to account for any overdispersion beyond that expected under a binomial or Poisson distribution.

As described above, assays of viability in outbred lines and assays of male mating success in both MA/control and outbred lines were each conducted in two blocks. We tested for a fixed effect of block in each case. Block did not have a significant effect on viability in outbred lines (*P* = 0.78), and was subsequently dropped from that model. Block had a significant effect on male mating success in outbred lines (*P* < 1 × 10^−5^), and controls (*P* < 0.05), and a marginally non-significant effect in MA lines (*P* = 0.06). However, we determined that there was no significant difference in genetic variance between blocks in any of these groups (5000 bootstrap replicates, all *P* > 0.35; see below for bootstrapping procedure details). We therefore modeled male mating success with a main effect of block on the intercept only.

We estimated 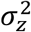 for each trait as the among line (genetic) variance among outbred lines on the log scale (see Supporting Information). We estimated *ΔM*_*z*_ as (E[*y*_*control*_] – E[*y*_*MA*_])/52, where *y* = log(*z*) and 52 is the number of MA generations. We also estimated the mutational variance in each trait (i.e., the per generation change in genetic variance of relative trait values among MA lines) as *ΔV* = *V*[*y*_*MA*_]/52. The “excess” variance in a trait beyond that expected under mutation selection balance alone was estimated as 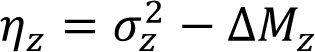. When comparing excess variance across traits we standardized ɳ_z_by the expected standing variance, Δ*M*_*z*_ (i.e., β_z_ = ɳ_z_/Δ*M*_*z*_). To attempt to account for selection on multiple traits, we *E*[*cz*] for each trait as 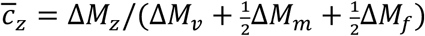 To attempt to account for selection on multiple traits, we estimated *E*[*c*_*z*_] for each trait as 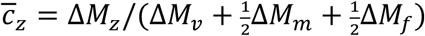, where the subscripts *v*, *m*, and *f* represent viability, male mating success, and fecundity, respectively. (The coefficients of ½ appear before *ΔM*_*m*_ and *ΔM*_*f*_ because these fitness components are sex limited.) We then estimated excess variance as 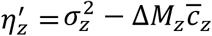.

We used bootstrapping to test hypotheses and obtain standard errors and confidence intervals for parameters of interest. In each of 20000 bootstrap replicates, line identifiers were sampled with replacement within each group (the two blocks for MA, control and outbred lines) and used to generate a dataset with the same structure as the actual data, including blocking. We then fit models to the resampled data to estimate means and genetic variances as described above. For variances, which cannot be negative, we report one-sided *P-* values and corresponding confidence intervals; all other tests are two-sided.

As an alternative analysis approach, we also fit generalized linear mixed models using the *R* package *MCMCglmm* (Hadfield 2010), using noninformative priors (variances ∼ 0, *nu* = –2), and a burn in phase of 10^6^ iterations. Subsequent iterations were stored such that the autocorrelation among stored values was less than 0.1 for all model parameters and the final number of stored iterations was 10^4^. To test hypotheses we compared the posterior distributions for the parameters of interest.

We also used *MCMCglmm* to examine the standing and mutational genetic correlations among traits. There is no between-trait association at the level of individual measurements in our data, and so we did not fit residual covariances. We selected priors that are expected to be uninformative for random effect correlations (inverse-Wishart; *G*: variances = 0.02, covariances = 0, *nu* = 4; *R*: variances ∼ 0, covariances = 0, *nu* = –2; see *MCMCglmm CourseNotes* vignette). Models were conducted with a burn-in phase of 10^6^ iterations followed by 10^8^ iterations with a thinning interval of 6000, leaving 16667 values in the posterior distribution such that autocorrelation among stored values was less than 0.1 for all model parameters. For each posterior sample, correlations between traits were determined after converting means, variances and covariances to the original scale by integration. We obtained credible intervals using *HPDinterval* in the *coda* package (Plummer et al. 2006). To compare mutational and standing correlations we sampled values from each posterior distribution 10^5^ times. In addition, we estimated phenotypic correlations among line means between pairs of traits, which will differ from the genetic correlations due to estimation error (attenuation). For male mating success line means were first standardized by subtracting the block mean. Phenotypic correlations were compared between groups by bootstrapping with 10^4^ replicates.

### Maximum likelihood

Inferences from the analyses above assume the ideal control. In reality, each of the three control populations for the MA experiment consisted of a population of N = 450 adults, constructed using the same isogenic chromosome used in the MA lines. Though purifying selection will prevent mutations from fixing in control populations, deleterious mutations will appear and segregate at a low level, reducing the average fitness of the controls below their initial value. The occurrence of deleterious mutations on the initially-isogenic control chromosomes would downwardly bias our estimates of *ΔM*. We make an attempt to account for this possibility using a more detailed, yet still simplistic, model described in the Supporting Information. After converting population-genetic parameters to trait means and variances, we evaluated the likelihood of a given set of parameters using MA, control, and standing variance data sets with the generalized linear mixed model functions from *lme4* as described above. We optimized the likelihood function using the Nelder-Mead algorithm with the *R* package *bbmle* (Bolker et al. 2017), repeating the optimization 50 times with random starting values. We obtained approximate 95% confidence limits for the excess variance factor *β* by optimizing at multiple fixed values of *β* to find the values producing a decrease in log-likelihood of 1.92 units.

To evaluate the statistical power of tests of the mutation-selection balance model we conducted approximately 40 000 simulations of mutation accumulation lines and outbred lines with known values for excess variance and variation in sample sizes and other parameters. Power was estimated as the proportion of simulations in which significant (P < 0.05) excess variance was detected when applying the bootstrapping procedure described above. Simulation results were analyzed with analysis of variance (see Supporting Information).

## RESULTS

Our goal was to test the hypothesis that genetic variance reflects mutation selection balance in a more quantitative fashion than has been attempted previously, and to do so for multiple fitness components. To do this we collected data on the heterozygous effects of mutant and outbred chromosomes for three fitness components (traits).

Means and variances for each trait on the original scale of measurement are given in Table S1, and estimates of quantitative genetic values are shown in Table 1. While there is considerable variation among previous studies, our estimates of mutational decline and standing variance are consistent with previously reported values (Houle 1998; Charlesworth and Hughes 2000; Halligan and Keightley 2009). Using bootstrapping, we detected significant mutational decline in male mating success (*ΔM*_*m*_ > 0: *P* < 0.05) and viability (*ΔM*_*v*_ > 0: *P* < 0.01), but not female fecundity (*P* = 0.12). There is evidence for significant mutational decline when combining evidence across traits (*Z* = –8.35, *P* < 0.0001). *ΔM* did not differ significantly among traits (all *P* > 0.093), and did not differ significantly between adult fitness (averaging across sexes) and viability (*P* = 0.45). We detected significant mutational variance for female fecundity (*ΔV*_*f*_ > 0: *P* < 0.05), but not for the other traits (*P* > 0.18 for both). There is evidence for significant mutational variance when combining evidence across traits (*Z* = –4.23, *P* < 0.05).

**TABLE 1.**
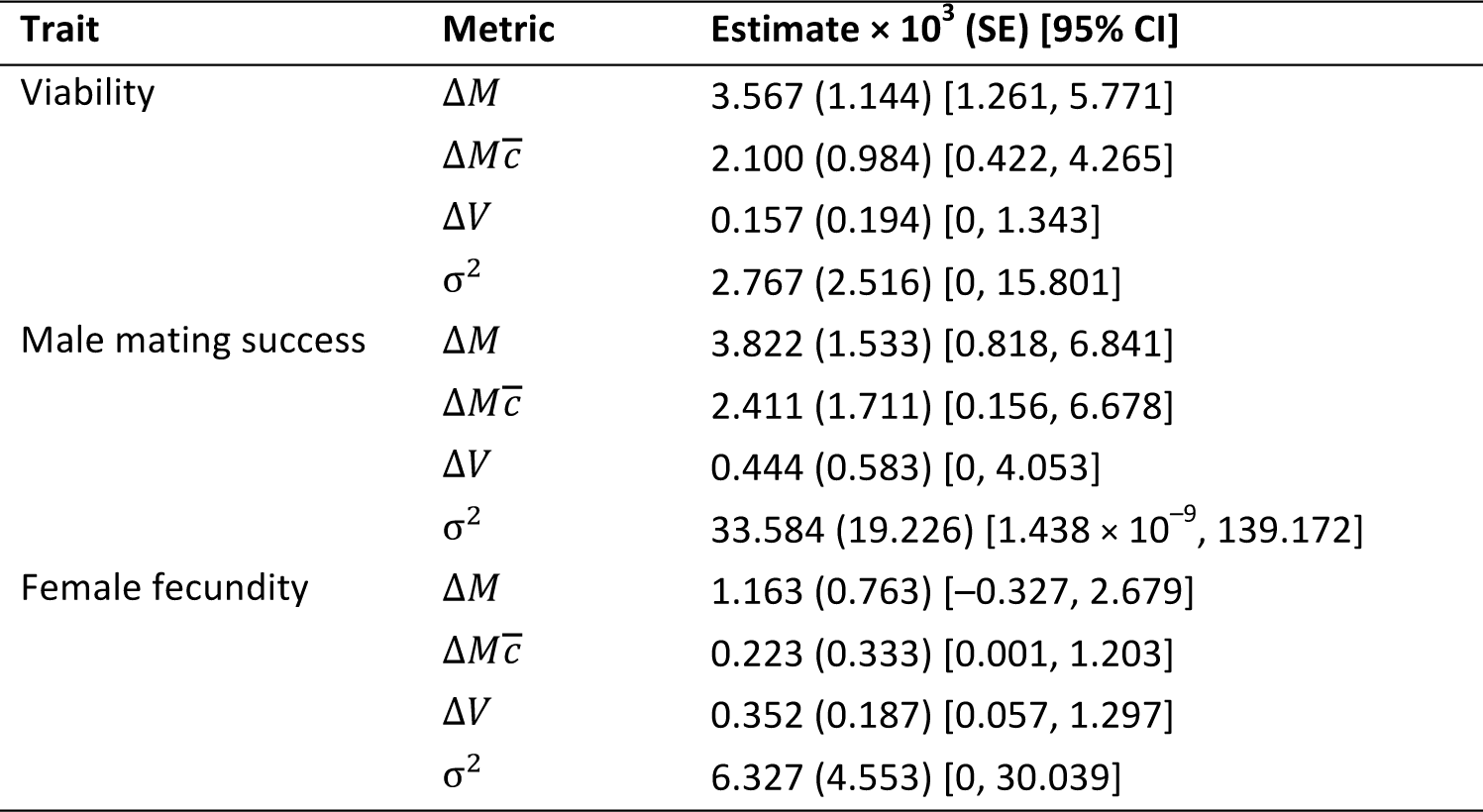
Summary of quantitative genetic estimates for each trait, for haploid second chromosomes

Our estimates of the standing genetic variance in relative trait values are associated with high uncertainty, but standing genetic variance was significant for male mating success 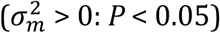 and marginally non-significant for female fecundity 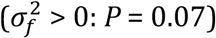 for viability *P* = 0.12. There is evidence for significant genetic variance when combining evidence across traits (*Z* = –5.19, *P* < 0.01).

Our primary interest was in whether mutation-selection balance can account for observed levels of standing variance. Based on point estimates, there was more standing variance in male mating success and female fecundity than expected under mutation-selection balance, whereas the opposite was true for viability (Figure 1). Our best point estimates of excess variance are as follows. For male mating success and female fecundity, respectively, there was 8.9 and 5.4 times more standing variance than expected under mutation-selection balance. In contrast, for viability the standing variance was 22% less than expected. However, there is high uncertainty in estimating excess variance. While the difference between observed and expected variance, 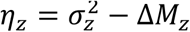, was not significant for any of the traits individually (*P* > 0.15 for all traits), the combined evidence from the adult fitness components indicates significant excess variance 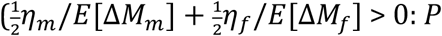, and excess variance in the adult fitness components was significantly greater than the excess variance for viability greater than the excess variance for viability 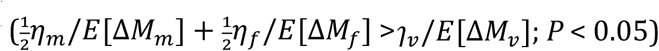. Using simulations, we assessed the statistical power of this approach to detect various levels of excess variance given our best estimates of the other relevant parameters (Supporting Information). These simulations indicate differences in detection power among traits (Fig. S4), and suggest that an experiment like ours would detect significant excess variance in fecundity or viability more than half the time when the observed variance exceeds expected variance by ∼5 times, whereas our power to detect the same degree of excess variance is much lower for male mating success.

**FIGURE 1.**
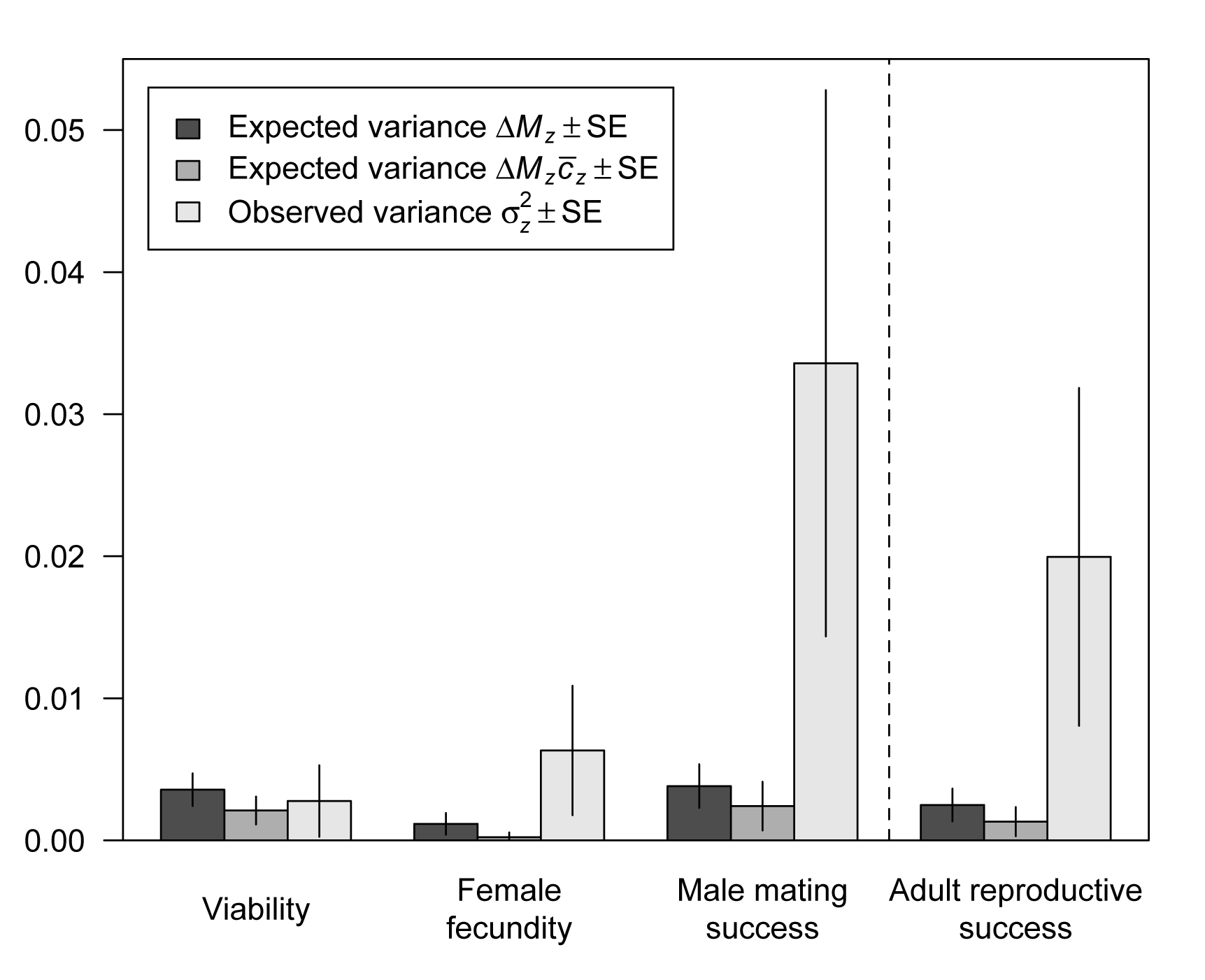
Genetic variance expected under mutation-selection balance and observed genetic variance for viability, male mating success, female fecundity, and sex-averaged adult reproductive success.

Our estimates of excess variance may be downwardly biased, because mutations will generally have greater effects on total fitness than their effects on any one fitness component 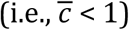 We can attempt to reduce this bias by accounting for 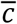 based on these three fitness components 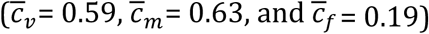, reducing the expected variance in all traits (Figure 1). Accounting for *c* does not qualitatively alter the statistical results described above; this is the case whether or not we exclude bootstrap replicates where one or more *ΔM*_*z*_ values is negative (6.7%), which results in nonsensical values for 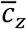. Accounting for 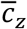, our best estimates are that there was ∼13.9 and ∼28.4 times more standing variance than expected under mutation selection balance for male mating success and female fecundity, respectively, whereas there was only ∼1.3 times more variance than expected for viability. Accounting for 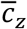, we find significant excess variance in sex averaged adult fitness (*P*:< 0.05;), with mutation-selection balance explaining only 6.6% of standing variation (Figure 1).

Results of the *MCMCglmm* models are very similar to the bootstrap results (Fig. S3 and Table S2). However, with these models we also find evidence for significant excess variance in male mating success (ɳ_m_<0:*P*<0.05;accounting for 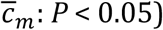), and female fecundity (ɳ_f_ <0:*P* =0.08 accounting for 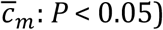). Here, our best estimates are that the standing variances are ∼11.2, 6.8, and 1.2 times larger than expected under mutation selection balance for male mating success, female fecundity and viability respectively (∼17.6, 35.6 and 2.1 when accounting for 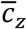).

To address the possibility that deleterious mutations may have reduced the fitness of control populations causing us to underestimate *ΔM*, we fit a maximum likelihood model designed to account for this effect. The full results of this model are shown in Table S3. As expected if deleterious mutations reduced fitness in the controls, we find that the ancestral trait values inferred by maximum likelihood exceed the observed control means, leading to higher estimates of *ΔM* under the maximum likelihood model. The maximum likelihood estimates for the ratio of standing genetic variance relative to that expected are associated with high uncertainty and are not significantly different from 1; mean [95% CI]: male mating success 3.6 [1, 11.5]; female fecundity 1.1 [1, 2.6]; viability ∼1 [1, 1.6] (note the model assumes that this ratio cannot be less than one). Thus, when we account for deleterious alleles segregating within the controls, we infer less excess standing variance but we still infer the same pattern among traits (i.e., *β_male_* > *β_female_* > *β_viability_*). This model finds evidence for significant mutational effects on all traits (likelihood ratio tests with 2 df comparing to model with *E*[*a*_*z*_] = *V*[*a*_*z*_] = 0: male mating success: *P* < 10^−3^; female fecundity: *P* < 10^−7^; viability: *P* < 10^−6^); this is perhaps unsurprising given that the model accounts for reductions in control fitness that would otherwise make mutational effects harder to detect.

The correlations between traits for both outbred lines and mutation accumulation lines are shown in Table 2. Relationships among line means are plotted in Figure 2. None of the genetic correlations are significantly different from zero, and there is no evidence of a difference between mutational and standing genetic correlations for any pair of traits (all *P* > 0.56). Although the phenotypic correlations based on line means are generally smaller than the genetic correlations (Table 2), we detect significant standing correlations between viability and female fecundity (*r* = 0.21, *t* = 2.41, *P* < 0.05), and between male mating success and female fecundity (*r* = 0.20, *t* = 2.30, *P* < 0.05). There is no evidence of a difference between mutational and standing phenotypic correlations for any pair of traits (bootstrapping; *P* > 0.40).

**TABLE 2.** Summary of genetic and phenotypic (line mean) correlation estimates for each pairwise trait combination, for mutation accumulation (MA) lines and the standing population.

| Group | Traits<sup>†</sup> | Genetic correlation (SE) [95%CI<sup>‡</sup>] | Line mean correlation (SE) [95%CI] |
| --- | --- | --- | --- |
| MA | m-f | 0.118 (0.321) [-0.479, 0.709] | 0.077 (0.146) [-0.211, 0.354] |
| | m-v | 0.117 (0.319) [-0.482, 0.7] | $2.94 \times 10^{-4}$ (0.148) [-0.278, 0.301] |
|  | f-v | 0.079 (0.274) [-0.441, 0.59] | -0.045 (0.149) [-0.339, 0.245] |
| Standing | m-f | 0.172 (0.25) [-0.318, 0.626] | 0.197 (0.073) [0.045, 0.328] |
|  | m-v | 0.182 (0.234) [-0.278, 0.615] | 0.096 (0.115) [-0.107, 0.329] |
|  | f-v | 0.278 (0.183) [-0.073, 0.628] | 0.206 (0.085) [0.036, 0.368] |

<sup>†</sup> Traits are viability (v), male mating success (m), and female fecundity (f).
<sup>‡</sup> Bayesian credible interval based on highest posterior density.

**FIGURE 2.**
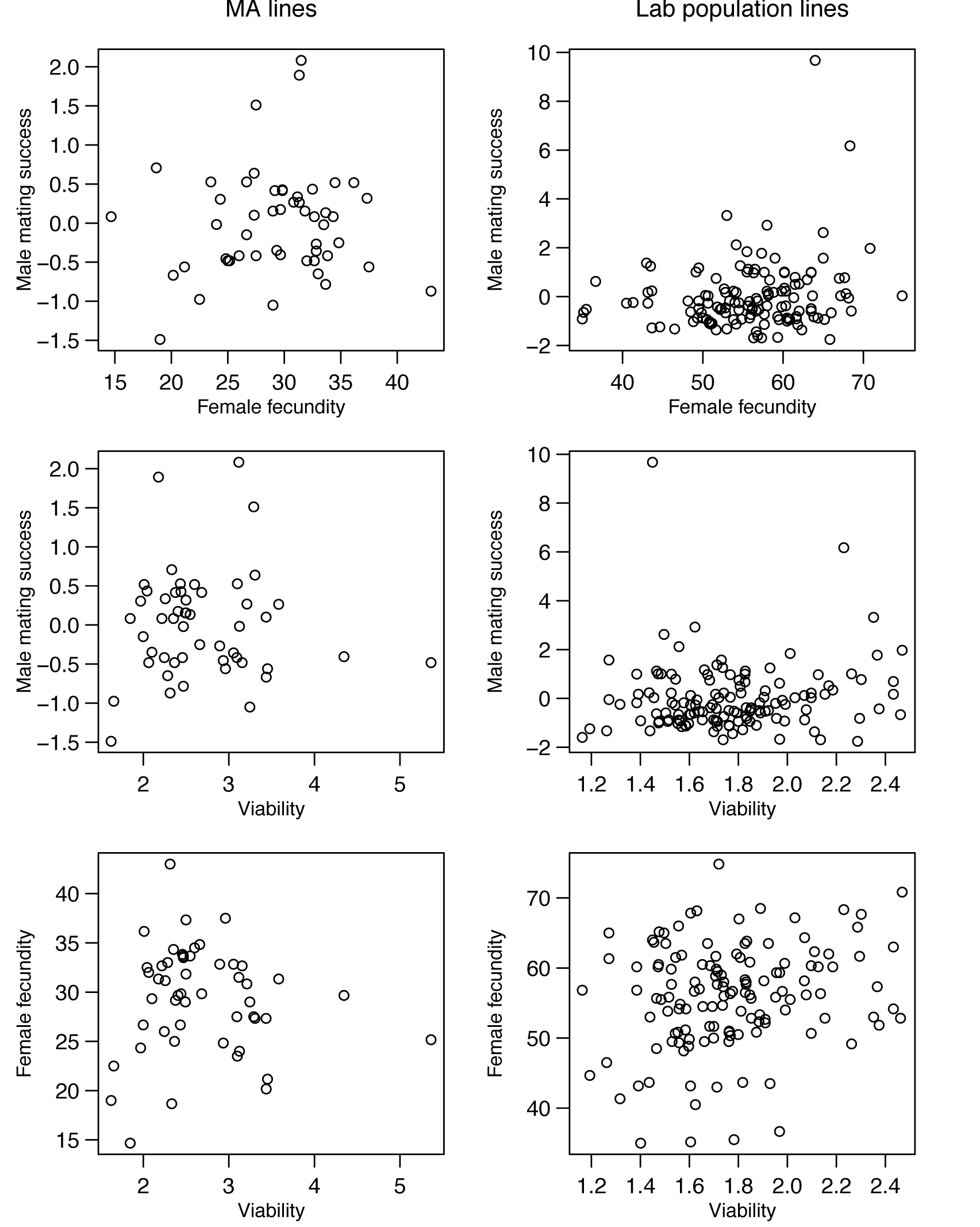
Phenotypic means for MA lines (left) and standing lab population lines (right), plotted for each pair of traits. Note that values for male mating success are standardized by the group mean within each block.

## DISCUSSION

We have attempted to test the hypothesis that genetic variance is maintained by mutation selection balance, and our results provide some support for additional sources of genetic variance. However, the fact that ancestral controls were able to evolve over the course of our experiment – which is an issue that affects many studies of mutation in *Drosophila* – could cause us to underestimate the contribution of mutation to standing genetic variance. When we attempt to account for this effect, we cannot reject a model where all standing genetic variance is due to mutation-selection balance.

One strength of our study is that we measured mutational decline and standing genetic variance for the same fitness components in the same genetic background. We are not aware of any other test of this kind. A partial exception is a study of the nematodes *Caenorhabditis elegans* and *C. briggsae* (Salomon et al. 2009), which compared mutational variance (i.e., ΔVz, not ΔMz) with standing genetic variance for lifetime fitness and body size across populations of each species. This ratio, 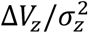, approximates the average strength of selection against heterozygous mutations, assuming mutation-selection balance (Barton 1990); given the values in Table 1, our estimates of 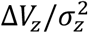 are 0.057, 0.013, and 0.056 for viability, male mating success, and female fecundity, respectively. These values are similar to other estimates based on the ratio 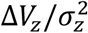 (Houle *et al.* 1996; Salomon *et al.* 2009), although they should be treated with caution since the denominator is not always significantly different from zero. However, if some standing variance is due to-non mutational sources then these values will be downwardly biased, and our results suggest that this could be the case for some traits, particularly male mating success. Huang et al. (2016) used a similar approach to compare mutational and standing variance in quantitative traits in *D. melanogaster* (e.g., bristle number), and concluded that simple models of mutation–stabilizing-selection-balance are insufficient to account for levels of standing variance.

Several others have made important contributions to this question by compiling relevant estimates from numerous sources, which are typically laboratory experiments using *D. melanogaster*. These authors compare several kinds of quantitative genetic estimates to assess the adequacy of the mutation-selection balance model and alternative models, but we will focus on their comparisons of mutational parameters and standing genetic variance because that was the approach taken in our study.

Houle et al. (1996) examined the ratio of standing variance to mutational variance, 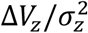, which approximates the “persistence time” of new mutations in a population under mutation-selection balance (Crow 1979). They reasoned that life history traits, which are under strong directional selection, should show reduced 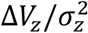 relative to metric traits, which tend to be under weaker selection. Balancing selection on alleles affecting life history traits would inflate the standing genetic variance, making the ratio more similar between trait classes. Consistent with mutation-selection balance, they found that the persistence time for life history traits (48) was significantly lower than that of metric traits (115). For comparison, our estimates of persistence time are 17, 76, and 18 for viability, male mating success and female fecundity, respectively (calculated using estimates from Table 1). Houle et al. (1996) discuss several challenges they faced, including the unavailability of estimates of 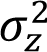 and ΔVz from the same population, possible bias resulting from using estimates based on homozygous rather than heterozygous mutations, and the use of standing variance measures from populations potentially not at equilibrium. Along similar lines, Houle (1998) found that 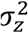 and ΔVz were highly correlated across traits, as expected if 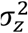 is largely determined by mutational input. However, the necessary standardization of traits may have introduced some autocorrelation to this relationship (Houle 1998).

One prediction considered by Charlesworth and Hughes (2000) was relationship 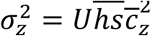 expected under mutation-selection balance (see their equation 19.6), which is equivalent to the relationship we tested (see Theoretical Background). They used the best available estimates of 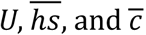 to calculate expected σ^2^ under mutation-selection balance. By comparison with observed values of σ^2^, they suggested that mutation likely contributes one-third to two thirds of the genetic variation in a typical life history trait. They did not attempt to determine the uncertainty surrounding this comparison, which is potentially quite high given the estimation error for each term. Nevertheless, our results are broadly consistent with their conclusion (average 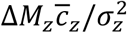 across traits = 0.29), although much lower for the adult traits (mean 0.05) than for viability (0.76). However, Charlesworth (2015) found, using a similar approach, that most *Drosophila* populations show higher standing variance for viability than expected under mutation-selection balance. This is not consistent with our finding for viability, which shows the least evidence for excess variance of the traits we studied, but again the statistical uncertainty surrounding the previous comparisons was not quantified.

A strength of the meta analyses described above (Houle et al. 1996, Houle 1998, Charlesworth and Hughes 2000, Charlesworth 2015) is that averaging over populations and traits may lead to more precise estimates of the parameters of interest. However, this approach might also obscure real variation among traits or populations in the extent of non-mutational genetic variance. Using a different method based on the effect of artificial selection on the mean value and inbreeding depression in a trait (Kelly 1999), there is evidence that intermediate-frequency alleles (i.e., not rare alleles at mutation-selection balance) contribute to variation in flower size and male fitness in *Mimulus guttatus* (Kelly and Willis 2001; Kelly 2003) and early fecundity in *D. melanogaster* (Charlesworth *et al.* 2007). Although it provides a qualitative test only, this approach is relatively assumption free, and it would be valuable to apply it to compare more traits and populations.

If some genetic variation is maintained by balancing selection, then balanced polymorphisms might be revealed by genome-wide scans of nucleotide variation. Although specific cases have been observed (e.g., related to self-incompatibility or immunity) this type of locus does not seem to be common, at least in model organisms (Tian *et al.* 2002; Charlesworth 2006) though there are challenges in detection. However, balanced polymorphisms may have disproportionately large effects on genetic variance, and so a small number of such loci could be sufficient to explain cases of non-mutational variation.

Additional evidence comes from the observation that mutation does not seem to completely explain the effects of inbreeding on life history traits (Charlesworth and Hughes 2000). Overall, these comparisons indicate that the mutation-selection balance model may often be insufficient to account for standing genetic variation. The data are inconsistent with heterozygote advantage across all traits, so any excess variance is likely due to balancing selection in the form of antagonistic pleiotropy (leading to net heterozygote advantage), heterogeneity in selection, or frequency-dependent selection (Charlesworth and Hughes 2000).

We tested the mutation-selection balance hypothesis by comparing mutational decline and standing variance in three fitness-related traits. We avoided assumptions regarding dominance by measuring the fitness effects of mutations in the heterozygous state. The overall effect of mutation accumulation was much weaker than the homozygous effects recorded previously for these lines (Sharp and Agrawal 2012, 2013), as new mutations are partially recessive (Simmons and Crow 1977; Phadnis & Fry 2005; Agrawal & Whitlock 2010; Manna et al 2011). Nevertheless, spontaneous mutations had significant effects on all three traits, in terms of either a decrease in the mean or an increase in the variance (Table 1). We measured standing genetic variation in each trait with the same procedure, using second chromosomes on the same isogenic background as the MA chromosomes, allowing us to statistically compare predicted and observed genetic variance in each trait.

For fitness itself, the ratio 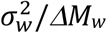 represents the standing variation in fitness relative to that expected from recurrent mutation. For a component of fitness, *ΔM* will arguably represent the maximum variance that can be due to mutation because the frequency of deleterious alleles will also be reduced via selection on additional components of fitness (assuming positive pleiotropy). For sexual traits this ratio was relatively high but was lower for larval viability (Figure 1). When we attempt to account for selection on the other traits, the ratios become smaller for all traits, but are still more extreme for the sexual traits. These results are consistent with the prediction that sexual traits may harbour more non - mutational variation than non-sexual traits.

If alleles with sexually antagonistic effects are responsible for the excess variance in sexual traits, then sexually antagonistic variation should also affect the standing genetic correlation between traits. A hallmark of intralocus sexual antagonism is a negative intersexual correlation for fitness (Chippindale *et al.* 2001). We found that the standing correlation between male mating success and female fecundity was positive and not significantly different from zero (Table 2). However, the standing correlation will also reflect the effects of deleterious alleles, which are thought to be positively-correlated among fitness components, including male and female reproductive success (Houle *et al.* 1994; Keightley and Ohnishi 1998; Mallet *et al.* 2011; Sharp and Agrawal 2013). Sexually antagonistic alleles should therefore generate a standing intersexual correlation that is not necessarily negative, but less positive than the correlation expected under mutation selection balance alone. However, we found neither statistical support for a negative standing genetic correlation (all point estimates are positive) nor statistical support for the standing genetic correlation being less than the mutational correlation (point estimates show the opposite pattern). However, our confidence in these negative results is low, due to the uncertainty in each estimate.

Another interpretation of our results is that some variance in male mating success and female fecundity may be due to non-sexually antagonistic balancing selection. Alleles affecting adult fitness, but not viability, could be involved in antagonistic pleiotropic interactions with fitness components not measured in this experiment (e.g., adult viability, sperm competitive ability), or be subject to some other form of balancing selection. Furthermore, although laboratory populations are maintained under relatively constant conditions, there may still be environmentally heterogeneous or fluctuating selection acting on these traits, or on correlated traits.

Another possible explanation arises from the fact that the conditions of the fitness assays do not represent selection on these fitness components exactly as it occurs in the laboratory population. For example, if the effects of mutations on a given fitness component had a larger effect in the assay conditions than in the laboratory population, this would inflate our estimates of mutational decline and standing variance, but would have a greater effect on standing variance, giving the false impression of excess variation. Our results could therefore be explained by our adult fitness assays being more selective than reality and our viability assays being less selective. As a first approximation, our fitness assays should resemble fitness as it occurs in the laboratory population but these assays are not identical to the conditions under which the population was maintained. Indeed, selection on both larval and adult fitness components are known to be somewhat sensitive to assay conditions (Kondrashov & Houle 1994; Zikovitz & Agrawal 2013). *Drosophila* populations with maintenance regimes that can be more precisely replicated in fitness assays would be preferable (e.g., Mallet et al 2011; Gibson et al 2002).

Although our data represent one of the more controlled tests to date of mutation selection balance in fitness components, several empirical limitations of our study should be considered. First, new mutations that arose in the control populations during the mutation accumulation procedure cause us to under-estimate mutational decline and over-estimate excess variance; we attempted to account for this in our maximum likelihood model but this is not an ideal solution. Mutational decline in control populations may be unavoidable in *D*. *melanogaster*, but can be prevented in model systems where ancestral genotypes can be cryopreserved.

Second, we chose to measure mutational decline and standing genetic variance in three traits at once, which placed a practical limit on the number of independent genotypes we could manipulate and examine. One of our goals was to make a legitimate attempt to quantify the statistical uncertainty in tests of mutation-selection balance. Testing multiple traits with only moderate replication contributed to that uncertainty, which is particularly problematic for traits where power is inherently limited (Fig. S4). Nevertheless, we gained some statistical power by combining evidence across traits.

Third, we measured standing variation on only an autosomal part of the genome. This was partly done for practical reasons: we wanted to compare standing variation with mutational variation, which arose on the second chromosome in our MA lines, and we wanted to avoid complications due to hemizygosity in males. However, we might have been more likely to detect a departure from mutation-selection balance if we had included variation on the sex chromosomes, which may be enriched for alleles with sexually antagonistic effects (Rice 1984; Gibson *et al.*2002).

Finally, we measured fitness decline by allowing mutations to accumulate on a particular second chromosome (albeit with variation on the remaining chromosomes) whereas we assessed standing variation across 133 second chromosomes. Given the evidence for considerable variation in mutation rates among genotypes (Sharp and Agrawal 2013; Schrider *et al.* 2013; Ness *et al*. 2015), we cannot be sure how closely our measured rates of mutational decline reflect the mutational process across genotypes in the laboratory population. Changes in the mutation rate during MA or directional epistasis could also cause bias in our estimates of mutational decline. However, all fitness effects were measured on a common genetic background.

Taken at face value, our experiment suggests that, within a given population, mutation selection balance may be an adequate explanation for much of the genetic variance in some fitness components but not others. Comparing fitness components in this respect may lead to greater insight into the sources and genetic basis of non-mutational variation

## Acknowledgements

We thank S. Otto, M. Whitlock, and members of the Otto lab for helpful discussion. This research was funded by the Natural Sciences and Engineering Research Council (Canada) to NPS (Vanier Graduate Scholarship) and AFA (Discovery Grant).

## SUPPORTING INFORMATION

### Testing mutation-selection balance using measures on a log scale, with multiplicative effects of mutations

Assuming deleterious mutations reduce the value of a life history trait in a multiplicative fashion, the value of trait *z* in a given MA line will be

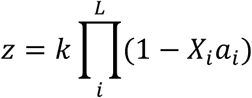

where *k* is the ancestral trait value, *L* is the number of loci at which mutations could occur, *X*_*i*_ is an indicator of the presence (1) or absence (0) of a mutation at locus *i*, and *a*_*i*_ is the heterozygous effect of a mutation at locus *i* on the trait.

It is useful to consider traits on the log scale, and we define

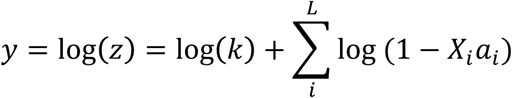

Note that using a Taylor series approximation shows that when *x* is small,

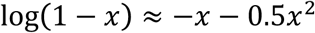

and we can write

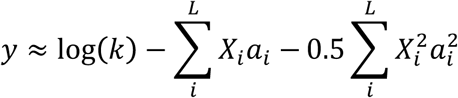

Assuming no covariance between the mutation rate and mutational effect at a given locus, the expected log trait value is

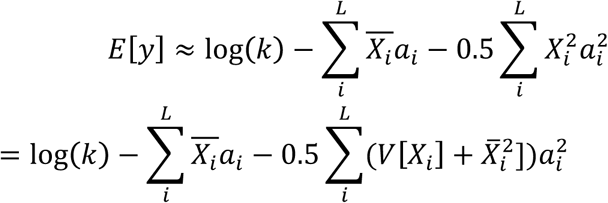

For the purposes of estimating *E*[*y*] we ignore terms in *a*^2^, which are small. Noting that *E*[*X*] summed over all loci can be written *Ut*, where *U* is the total rate of mutation across all loci and *t* is the number of generations of MA, we have

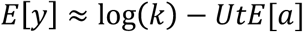

When the trait value of controls represents the ancestral trait value (which is likely not the case in our study, but applies to systems where static controls are available), the rate of change in the trait value per generation, *ΔM*, can be estimated as

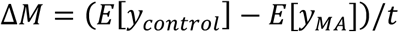

An approximation for the control mean in our experiment can be found below. To find the variance in trait values we need approximations for *E*[*y*^2^] and *E*[*y*]^2^.

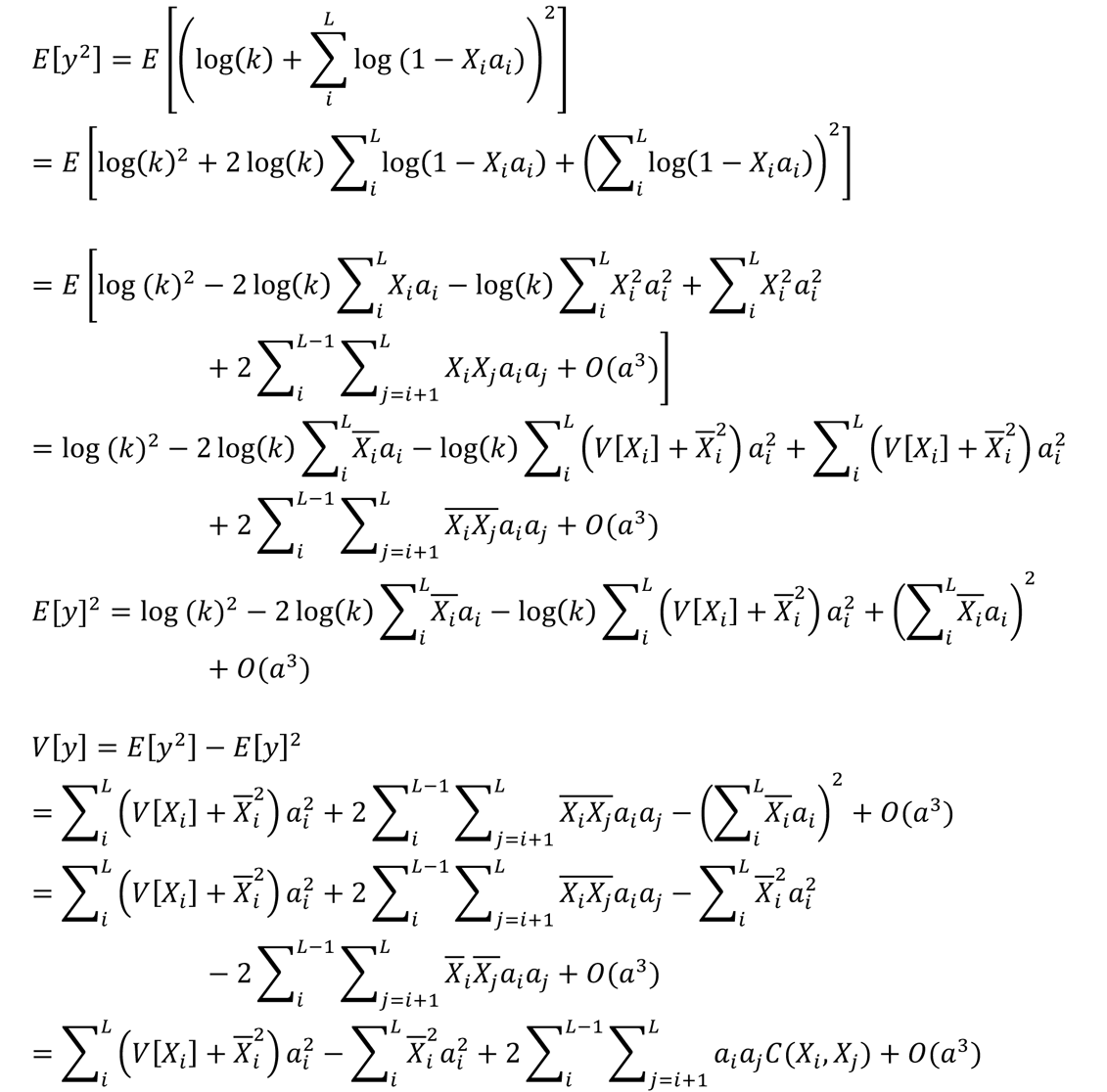

Assuming loci mutate independently, *C*(*X*_*i*_, *X*_*j*_) = 0, leaving

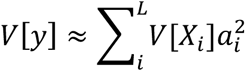

Assuming mutations arise at random among lines, i.e., following a Poisson distribution where *V*[*X*] = *E*[*X*], we can write

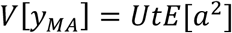

The change in trait variance per generation can therefore be estimated as

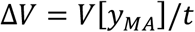

When the variance in a standing population at equilibrium is due only to segregating deleterious mutations, the above approach can be applied analogously, where *V*[*X*] = 2*pq*, giving

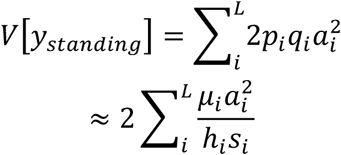

Because *a* = *chs*, we can write

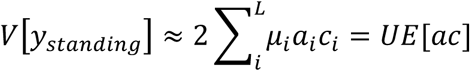

This relationship can also be found by considering standing variance in the relative untransformed trait value *z*

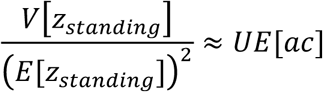

and noting that 9[log (2)] ≈ 9[2] ⁄ 8[2] ^7^ in a Taylor-series approximation.

### Maximum likelihood model

This model was implemented on the log scale, where *y* = log(*z*). Building from the material presented in the main text, we first describe how observed means and variances are related to underlying parameters. Genetic means and variances of MA lines:

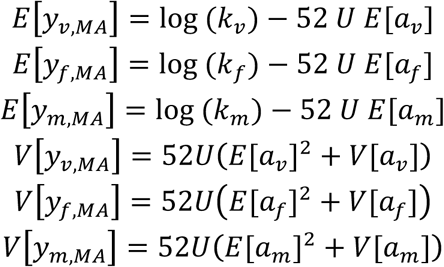

Mean and variance of selection on heterozygous mutations:

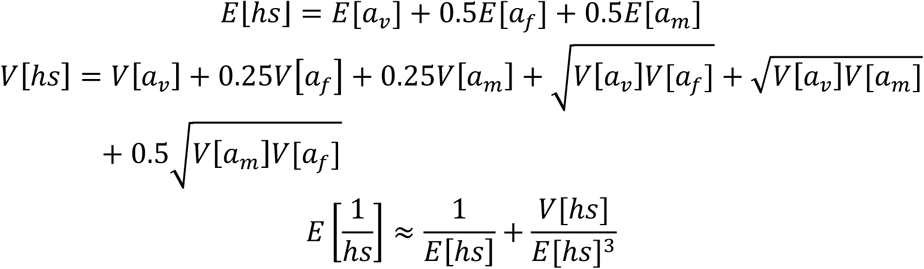

Standing genetic variance:

We can expand the standing variance predicted in trait *z* under mutation-selection balance as

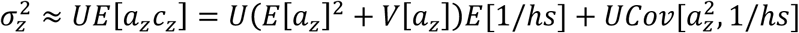

The last term above should be negative when pleiotropy is positive, and not too large; we omit it from our model, which should be a conservative assumption with respect to the main test. Finally, we use the Taylor series approximation

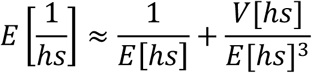

Allowing for variance from sources other than mutation selection balance, we model the standing genetic variance in the traits as

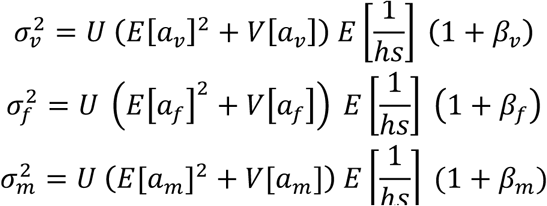

where β_2_ is the relative extent to which standing variance in trait *z* exceeds thewhere d_a_ is the relative extent to which standing variance in trait *z* exceeds the expected value, i.eβ ≡ (observed variance ⁄ predicted variance) –1. (d is equivalent to excess variance η divided by predicted variance).

Trait means of the control lines for the MA experiment:

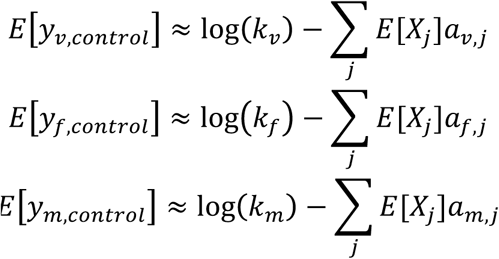

where *E*[*X*_*j*_] is the expected number of alleles on a control chromosome in deleterious size class *j* (defining the original copy of chromosome 2 used in this experiment as wild-type at all sites).

Ideally, the control would be mutation free (i.e., an exact replica of the original copy of chromosome 2, so that E[*X*_*j*_] = 0 for all *j*). In reality, deleterious alleles will segregate at low frequencies within the control populations so E[*X*_*j*_] > 0. Under a deterministic model, the average number of mutations of size class *j* in the controls after *t* (=52) generations is

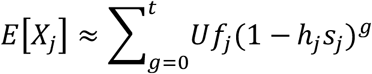

where *f*_*j*_ is the fraction of all new mutations that are in class *j*. We assume mutational fitness effects follow a gamma distribution (Keightley 1994) with mean *E*[*hs*] and variance *V*[*hs*] (shape *E*[*hs*]^2^/*V*[*hs*] and scale *V*[*hs*]/*E*[*hs*]), and divide the distribution into 100 discrete size classes, which we use to assign values to *f*_*j*_ and *h*_*j*_*s*_*j*_. We assume the effect of a mutation on total fitness can be approximated by its effect on the fitness components we studied, such that E[*hs*] = *E*[*a*_*v*_] + E[*a*_*f*_] /2 + E[*a*_*m*_] /2 and

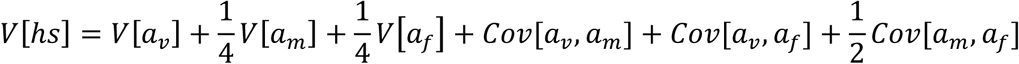

We assume the traits are perfectly positively correlated such that *Cov*[*a*_1_, *a*_2_] =*SD*[*a*_1_]*SD*[*a*_2_] because this is conservative with respect to the test for excess genetic variance.

Similarly, we can model the genetic variance of control lines assuming that the number of mutations of a given effect size class is Poisson-distributed among lines, such that *V*[*X*_*j*_] = *E*[*X*_*j*_]:

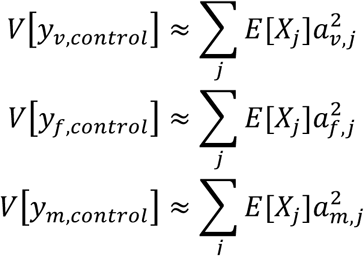

The remaining parameters are not of primary interest, and are free to vary, namely the mean trait values for outbred lines (3 parameters), block effects on male mating success (3 parameters) and overdispersion for female fecundity and larval viability (6 parameters). The 25 total parameters are listed in Table S3.

There are nine ‘sets’ of data (control, MA, and standing for each of the three traits). For each set we evaluate the log-likelihood of the data for given values of the mean, genetic variance, and overdispersion using the *lme4* generalized linear mixed model (*glmm*) deviance evaluation function for that subset. For male sets there is also a block effect on the mean, and overdispersion is absent. The overall log - likelihood is then the sum of the log-likelihoods across the nine sets. Those means and variances that are defined in terms of population-genetic parameters as described above. For example, the log-likelihood of the standing female fecundity set of data would be obtained by evaluating the *glmm* function using a specified set of values for an overdispersion parameter, the standing mean, and the standing variance, where the standing variance is given as a function of from underlying parameters 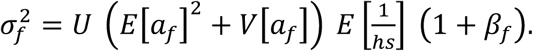

We evaluated the likelihood of a given set of parameters using the generalized linear mixed model functions from *lme4* as described above. We optimized the likelihood function using the Nelder-Mead algorithm with the *R* package *bbmle* (Bolker et al. 2017), repeating the optimization 50 times with random starting values. We obtained approximate 95% confidence limits for the excess variance factor *β* by optimizing at multiple fixed values of *β* to find the values producing a decrease in log-likelihood of 1.92 units.

### Power analysis simulations

#### Methods

To examine the statistical power of tests for excess variance we simulated data structures similar to those of our actual observations, on the log scale. We assessed power for three “traits” resembling female fecundity (counts with control⁄outbred mean *μ* = 3.7), male mating success (proportions with control⁄outbred mean *μ* = 0.7 and 6 trials per replicate), and larval viability (proportions with control⁄outbred mean *μ* = 0.9 and 50 trials per replicate). Control lines had zero genetic variance. MA line means were set to *μ – ΔMt*, with *ΔM* = 4 × 10^−3^ and *t* = 50 generations of MA. MA line genetic variances were set to *ΔVt* with *ΔV* = 2 × 10^−4^. Outbred lines had mean *μ* and genetic variance *ΔM*(1+ *β*), where 1+ *β* is the factor by which standing variance exceeds that expected under mutation selection balance. We evaluated *β* values of 1, 3, 7 and 11. In some simulations we allowed within line variance to exceed that expected under the Poisson or binomial process (overdispersion). When present overdispersion was set to 0.1 on the observation (non log) scale. The number of MA+ control lines was either 100 or 200, evenly split between MA and control. The number of outbred lines was either 100 or 200. The number of replicate measures of each MA or control line was set to 6. The number of replicate measures of each outbred line was set to 6, or 12 in some instances. Each parameter combination was simulated 400 times, and the test for excess variance was performed on each simulated dataset by bootstrap resampling among lines with 1000 replicates. This is fewer bootstrap replicates than our main analysis, but tests with higher bootstrap replication gave essentially the same results.

#### Results

Estimates of statistical power to detect excess standing genetic variance based on simulated data are shown in Figure S4. Power unsurprisingly increases with the magnitude of excess variance (*F* = 130.3, df = 1, *P* < 10^−15^), and also depends on the nature of the trait being examined (*F* = 107.7, df = 2, *P* < 10–15), with proportion data and counts with a relatively high number of trials per replicate (as in our measures of viability and female fecundity) providing greater power than proportion data with few trials per replicate (as in our measures of male mating success). The number of lines used to estimate standing variance had an impact on power (*F* = 7.8, df = 1, *P* <0.01; solid vs. dashed lines in Fig. 3), as did overdispersion (*F* = 9.5, df = 1, *P* < 0.01; panels A and B vs. C and D in Fig. 3). Over the range considered, the number of MA and control lines had no detectable influence on power (*F* = 0.001, df= 1, *P* = 0.98; panels A and C vs. B and D in Fig. 3). Considering the traits together, there was no evidence that allocating effort towards independent outbred lines versus replicate sublines had any effect on power (200 lines with 6 replicates each vs. 100 lines with 12 replicates each; proportion test: χ^2^ = 0.002, df = 1, *P* = 0.97).

**Table S1.** Means and genetic variances for each trait on the original scale of measurement

| Trait | Group | Mean (SE) [95% CI] | Genetic variance (SE) [95% CI] |
| --- | --- | --- | --- |
| Viability | Mutant | 2.721 (0.111) [2.530, 2.962] | 0.061 (0.081) [0, 0.671] |
| | Control | 3.302 (0.146) [3.030, 3.603] | 0.269 (0.172) [$3.138 \times 10^{-8}$, 1.122] |
|  | Standing | 1.758 (0.024) [1.712, 1.805] | 0.009 (0.008) [0, 0.050] |
| Male mating success | Mutant | 1.575 (0.083) [1.415, 1.736] | 0.059 (0.079) [0, 0.621] |
|  | Control | 1.951 (0.114) [1.717, 2.165] | 0.173 (0.138) [0, 1.103] |
| | Standing | 2.052 (0.083) [1.866, 2.191] | 0.148 (0.088) [$9.246 \times 10^{-9}$, 0.744] |
| Female fecundity | Mutant | 28.109 (0.733) [26.658, 29.519] | 14.587 (7.635) [2.362, 51.030] |
|  | Control | 29.948 (0.861) [28.249, 31.608] | 21.906 (7.546) [9.396, 53.537] |
|  | Standing | 53.052 (1.090) [50.784, 55.042] | 17.864 (12.654) [0, 80.040] |

**TABLE S2.** Summary of quantitative genetic estimates for each trait, for haploid second chromosomes, based on MCMCglmm models

| Trait | Metric | Estimate $\times 10^3$ (SE) [95% CI]<sup>†</sup> |
| --- | --- | --- |
| Viability | $\Delta M$ | 3.598 (1.113) [1.466, 5.822] |
| | $\Delta M\bar{c}$ | 2.111 (0.984) [0.422, 4.265] |
| | $\Delta V$ | 0.332 (0.242) [$1.900 \times 10^{-5}$, 0.797] |
| | $\sigma^2$ | 4.381 (2.897) [0.001, 9.735] |
| Male mating success | $\Delta M$ | 3.897 (1.697) [0.549, 7.177] |
| | $\Delta M\bar{c}$ | 2.476 (1.711) [0.156, 6.678] |
| | $\Delta V$ | 0.988 (0.716) [0.001, 2.354] |
| | $\sigma^2$ | 43.507 (24.198) [0.029, 87.232] |
| Female fecundity | $\Delta M$ | 1.172 (0.790) [-0.403, 2.692] |
| | $\Delta M\bar{c}$ | 0.224 (0.333) [0.001, 1.203] |
| | $\Delta V$ | 0.435 (0.188) [0.098, 0.805] |
| | $\sigma^2$ | 7.976 (4.440) [0.010, 15.940] |

<sup>†</sup> Bayesian credible interval based on highest posterior density.

**TABLE S3.** Summary of maximum likelihood model results. See Appendix for model parameterization.

| Parameter | Estimate | SE† |
| --- | --- | --- |
| $\log(k_m)$ | 1.102 | 0.164 |
| $\log(k_f)$ | 3.854 | 0.165 |
| $\log(k_v)$ | 1.603 | 0.157 |
| $E[a_m]$ | 0.024 | 0.005 |
| $E[a_f]$ | 0.019 | 0.002 |
| $E[a_v]$ | 0.021 | 0.004 |
| $V[a_m]$ | $1.023 \times 10^{-4}$ | $3.201 \times 10^{-4}$ |
| $V[a_f]$ | $1.678 \times 10^{-4}$ | $2.581 \times 10^{-4}$ |
| $V[a_v]$ | $3.587 \times 10^{-6}$ | not available |
| $U$ | 0.539 | 0.212 |
| $\beta_m$ | 2.600 | 0.003 |
| $\beta_f$ | 0.092 | 0.022 |
| $\beta_v$ | 1.615E-04 | 0.172 |
| $E[y_{m, standing}]$ | 0.687 | 0.038 |
| $E[y_{f, standing}]$ | 3.968 | 0.016 |
| $E[y_{v, standing}]$ | 0.563 | 0.014 |
| $\text{block}_{m, MA}$ | 0.106 | 0.055 |
| $\text{block}_{m, control}$ | 0.151 | 0.054 |
| $\text{block}_{m, standing}$ | 0.176 | 0.037 |
| $\text{error}_{f, MA}$ | 0.092 | 0.012 |
| $\text{error}_{f, control}$ | 0.073 | 0.010 |
| $\text{error}_{f, standing}$ | 0.141 | 0.011 |
| $\text{error}_{v, MA}$ | 0.117 | 0.024 |
| $\text{error}_{v, control}$ | 0.209 | 0.034 |
| $\text{error}_{v, standing}$ | 0.039 | 0.006 |

† Approximate standard errors based on quadratic approximation to the curvature at the maximum likelihood estimate. These values were not used for hypothesis testing in this study.

**Figure S1.**
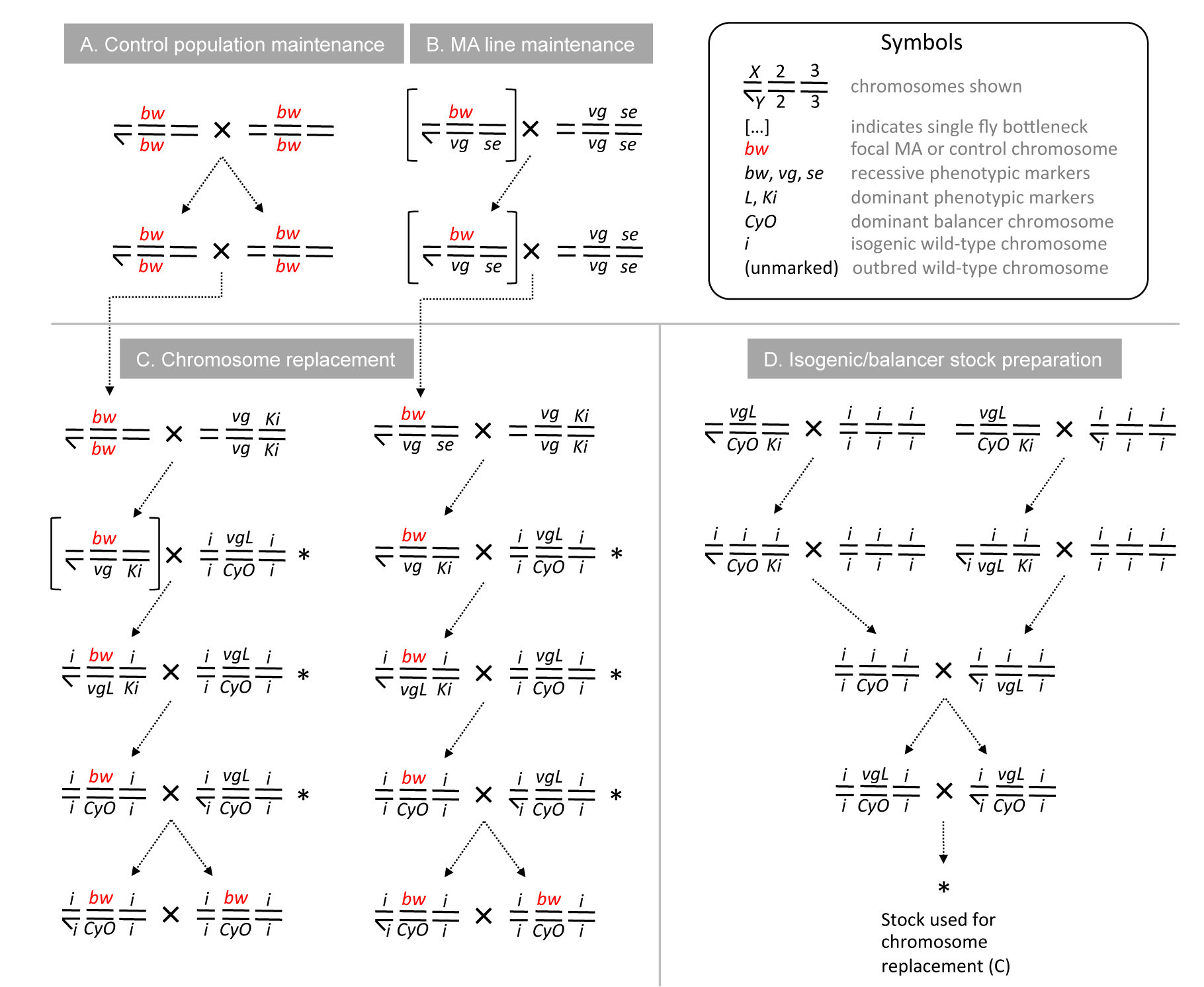
Details of mutation accumulation and control line maintenance and preparation. The first three chromosomes are shown for each genotype; the tiny fourth chromosome was not manipulated. Males generally lack recombination, and are identified here by the presence of a *Y* chromosome. Crosses took place using virgin females where appropriate. Chromosomes were identified using recessive phenotypic markers (*bw*, *vg*, *se*), dominant phenotypic markers (*L*, *Ki*), and a balancer chromosome (*CyO*), which suppresses recombination on the second chromosome. For mutation accumulation (MA), a single second chromosome marked with *bw* was used to initiate three control populations and numerous MA lines. These focal chromosomes are shown in red. (A) Control populations homozygous for the focal chromosome were maintained at a moderate size (450 adults) to prevent mutation accumulation. (B) MA chromosomes were propagated by bottlenecking to a single heterozygous male each generation, allowing new mutations to accumulate. (C) Following 52 generations of MA, crosses were performed to replace all non focal chromosomes with an isogenic background. Within line variation on the focal chromosome was eliminated by bottlenecking (square brackets). Each cross included 1-4 males and 4 females per line. These crosses involved several marker stocks, which were created using standard crossing methods (not shown). An isogenic stock with *vgL*⁄*CyO* was created as shown in (D), after creating a completely isogenic genotype using standard balancer chromosome methods (not shown).

**Figure S2.**
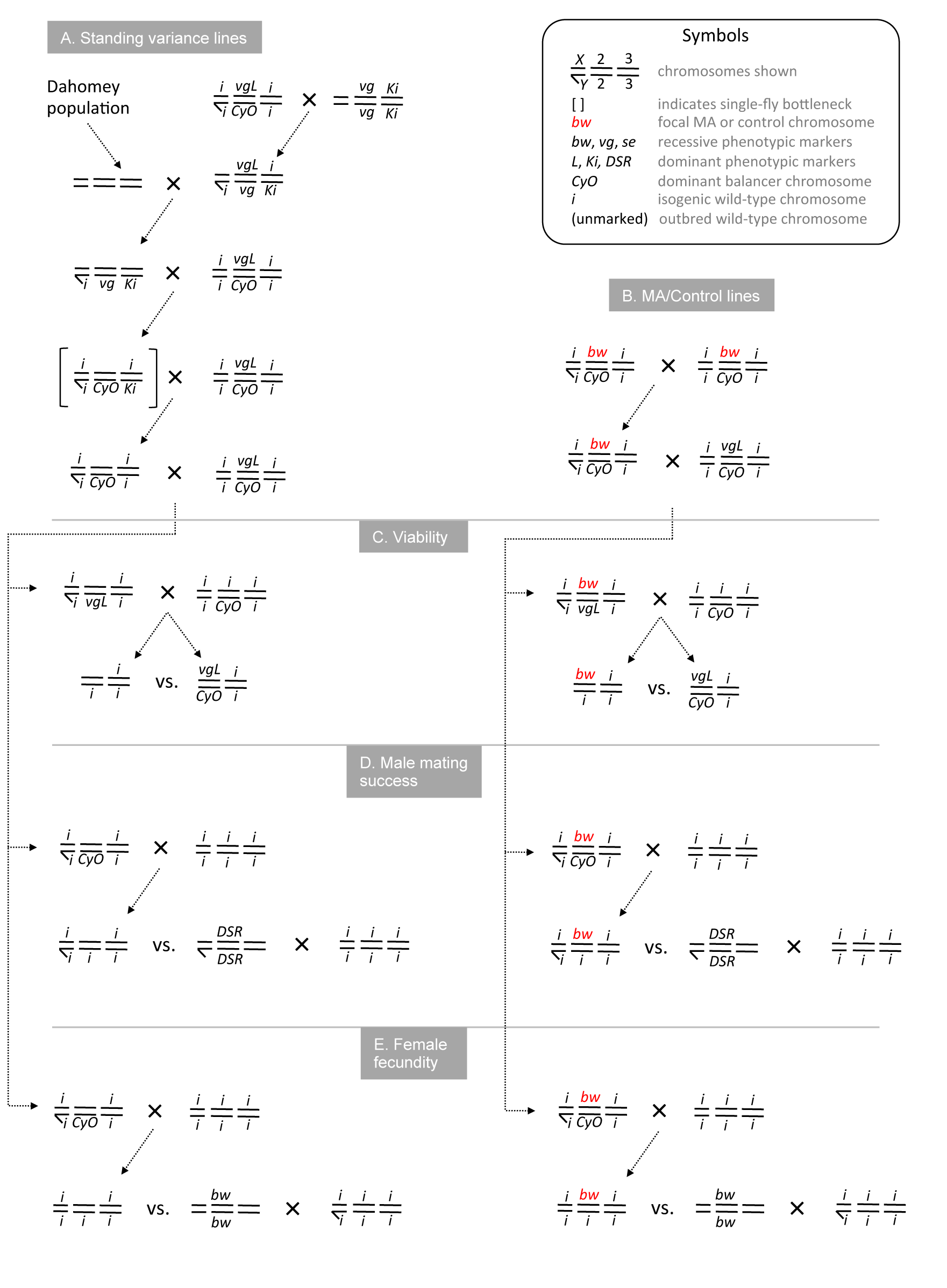
Details of standing variance line preparation and assay crosses for all lines. The first three chromosomes are shown for each genotype; the tiny fourth chromosome was not manipulated. Males lack recombination, and are identified here by the presence of a *Y* chromosome. Crosses took place using virgin females where appropriate. Chromosomes were identified using recessive phenotypic markers (*bw*, *vg*, *se*), dominant phenotypic markers (*L*, *Ki*, *DSR*), and a balancer chromosome (*CyO*), which suppresses recombination on the second chromosome. After obtaining focal second chromosomes from either the outbred Dahomey laboratory population (A) or MA lines and control lines (B; generated as shown in Figure S1), crosses were performed to assess viability (C), male mating success (D), and female fecundity (E), as described in the text.

**Figure S3.**
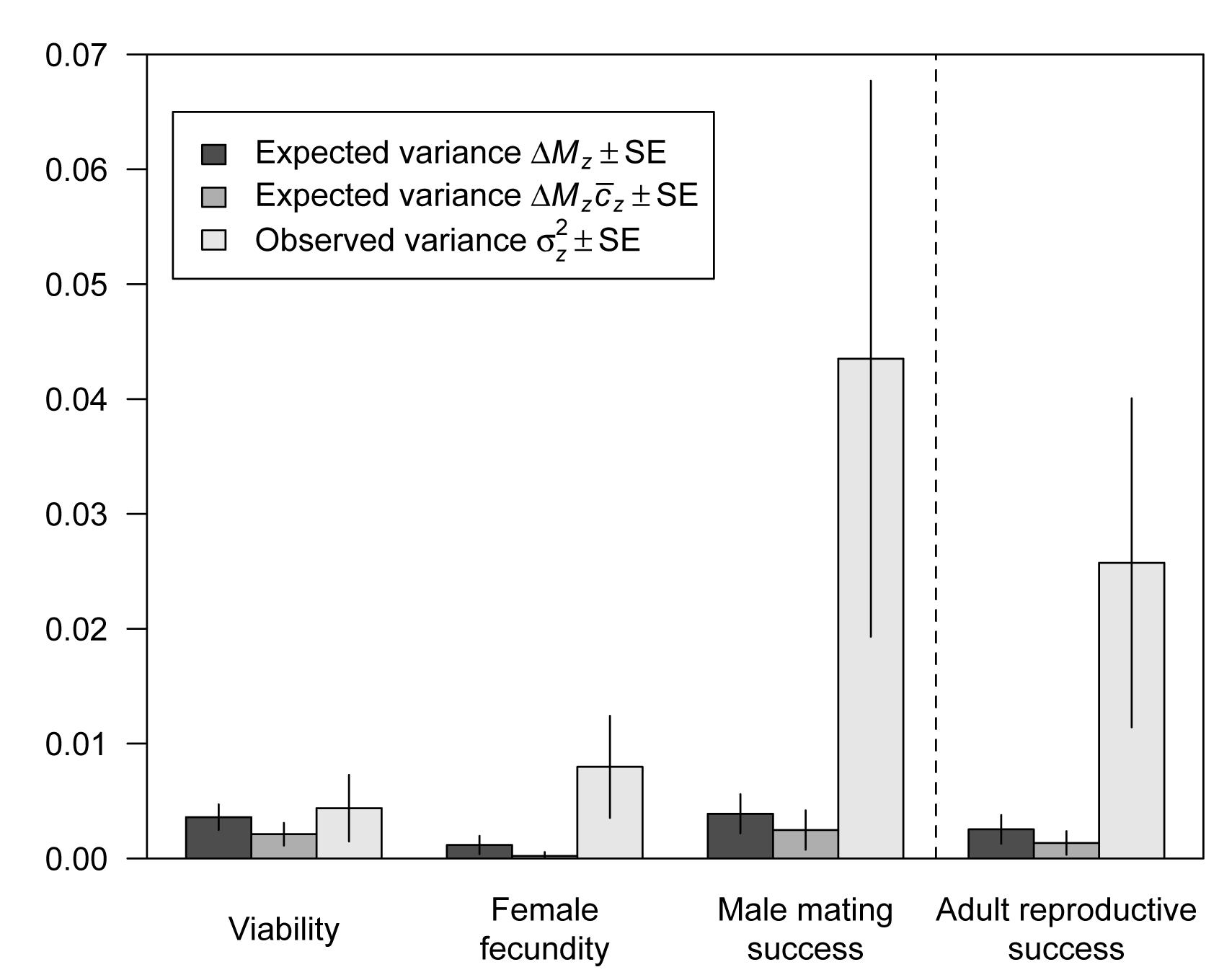
MCMCglmm model results. Genetic variance expected under mutation selection balance and observed genetic variance for viability, male mating success, female fecundity, and sex-averaged adult reproductive success.

**Figure S4.**
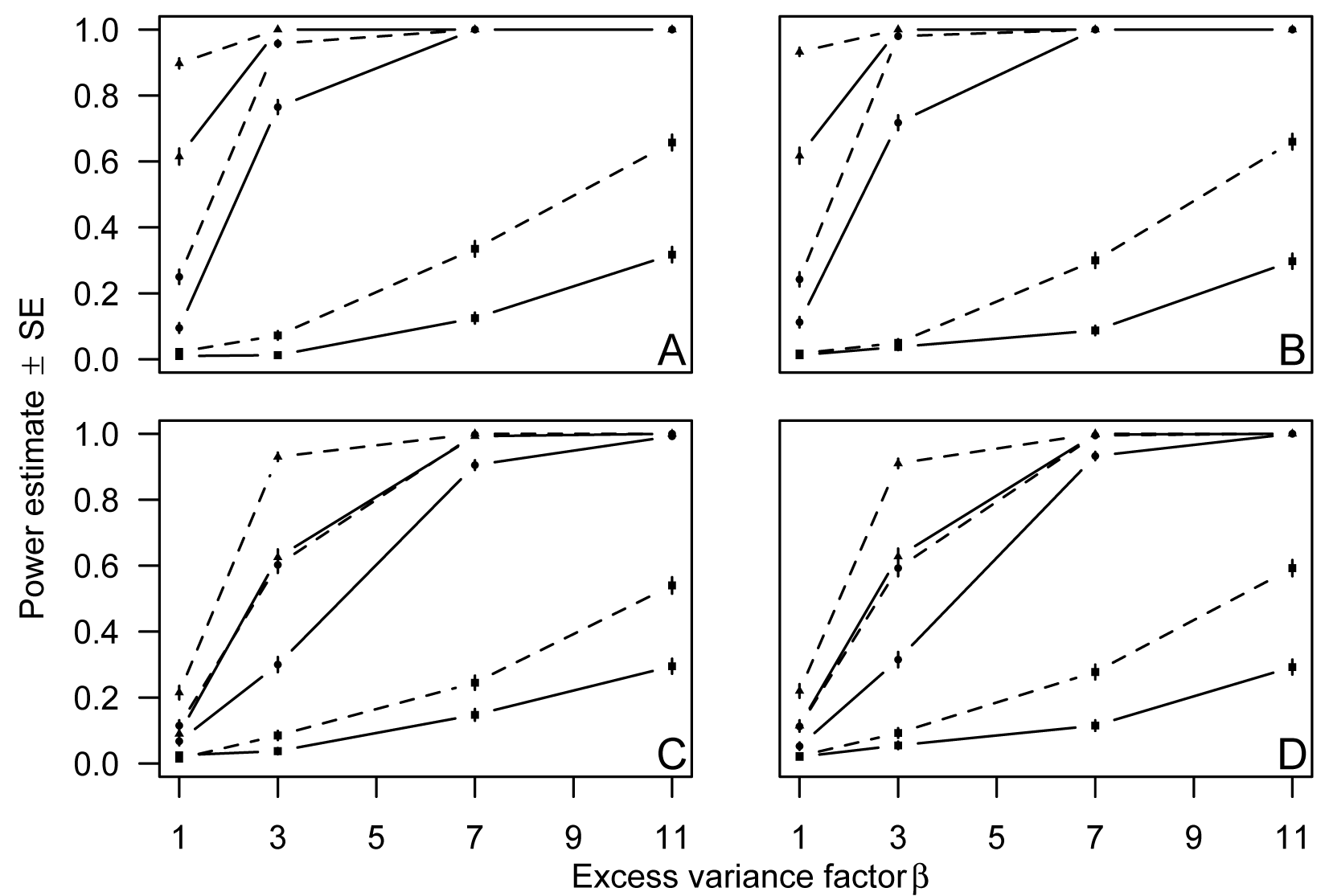
Power analysis results. Estimates of power to detect a significant excess of standing genetic variance compared to the expectation under mutation selection balance. Power was examined for three traits with structure resembling our measures of viability (circles; proportion with many trials), male mating success (squares; proportion with few trials), and female fecundity (triangles; counts with many events). The number of lines used to estimate standing variance was 100 (solid lines) or 200 (dashed lines). The number of MA and control lines used to estimate mutational decline was 100 (A and C) or 200 (B and D). Overdispersion was absent (A and B) or set to 0.1 (C and D).

